# Strain-level translocation and enrichment mechanisms of oral bacteria in the lower gastrointestinal tract of stunted children

**DOI:** 10.1101/2025.11.11.687814

**Authors:** Simon Yersin, Jean-Chrysostome Gody, Florent Mazel, Edgar Djimbele, Synthia Nazita Nigateloum, Bolmbaye Privat Gondje, Sonia Sandrine Vondo, Kaleb Kandou, Aline Raub, Youzheng Teo, Serge Ghislain Djorie, Nathalie Kapel, Philippe J. Sansonetti, Pascale Vonaesch, the Afribiota Investigators

**Author notes:** Corresponding author: Pascale Vonaesch. See in Appendix A (the Afribiota Investigators).

## Abstract

Emerging evidence suggests that ectopic colonization of oral bacteria in the lower digestive tract may exacerbate gastrointestinal disorders. Nevertheless, it remains unclear whether bacteria of oral origin are continuously translocating from the oral cavity to the lower gastrointestinal tract or are locally adapted and persist in their respective niches. We investigated strain translocation dynamics in 44 healthy and stunted children from Bangui, Central African Republic. Using cross-sectional shotgun metagenomic sequencing of saliva, gastric, duodenal and fecal samples, and isolation and whole genome sequencing of 87 *Streptococcus salivarius* isolates, we showed translocation of members of the genera *Streptococcus*, *Veillonella*, *Rothia* and *Haemophilus*. Fecal isolates were more closely related to oral isolates from the same individuals than those from other individuals. Additionally, saliva showed higher *S. salivarius* nucleotide diversity compared to other compartments, suggesting a source-sink dynamic in which *S. salivarius* populations are continuously seeded from the oral cavity without durably establishing in the lower gastrointestinal tract. Last, we showed that overrepresentation of oral bacteria in the duodenum of stunted children is due to increased biomass, while in the colon it is linked to depletion of overall biomass, including in butyrate-producing strains. Our study quantifies mechanisms of oral-to-gut translocation and enrichment of oral taxa, providing key insights into microbiota disruption in stunted children.

## Introduction

The human digestive tract is colonized by diverse microorganisms, including bacteria, viruses and fungi, that play a central role in both health and disease^1^. The oral cavity, small intestine and colon are host to bacterial communities distinct in their composition and function, which reflect the local adaptation of bacteria to their ecological niches within each body compartment^2–5^. Nevertheless, the digestive tract is connected by a constant flow of food and saliva which transports bacterial cells from the oral cavity to the lower gastrointestinal (GI) tract^6^. Although gastric acidity and antimicrobial compounds reduce the number of viable cells reaching the lower GI tract by several orders of magnitude, bacteria of oral origin are detected in the feces^6,7^. Accumulating evidence suggests that ectopic colonization by bacteria from the oral cavity exacerbate various GI and other inflammatory diseases^8^.

Stunted growth, defined as children that are more than two standard deviations below the height-for-age z-score (HAZ) of the WHO Growth Reference Cohort, affects more than 150 millions children under the age of 5 years^9^. Stunting is associated with small intestinal oral bacterial overgrowth (SIOBO, >=10^5^ CFU/mL), leading to inflammation and lipid malabsorption in the small intestinal tract as well as neurodevelopmental changes in a murine model of childhood undernutrition^10–13^. Most importantly, the absolute abundance of oral bacteria in the duodenum of stunted children has been correlated with stunting severity^14^. In the lower GI tract, stunting has been associated with a reduction in anaerobic butyrate-producing bacteria and an overrepresentation of bacteria of oral origin^10,15^. Together, these findings suggest that translocation and overgrowth (>=10^5^ CFU/mL) of oropharyngeal taxa in the lower digestive tract may play a significant role in the development of the pathophysiology of stunting^16,17^.

Currently, the extent and mechanism of bacterial translocation from the oral cavity to the lower GI tract remains poorly characterized. Schmidt et al. demonstrated that translocation and colonization of oral bacteria in the colon is both common and extensive in healthy adults^7^. However, the translocation mechanisms explaining the presence of species of oral origin in the lower GI tract remain poorly understood. There are two scenarios (Figure 1A) in which bacteria of oral origin can establish in the small intestinal tract: 1) the oral cavity may serve as a reservoir from which bacterial strains are continuously translocated resulting in a connected population along the digestive tract^7,18^ or 2) bacterial strains of oral origin may be locally adapted to their respective niches for long term residency, leading to compartment-specific populations that are largely segregated between the oral cavity, small intestine and colon^19,20^. While these two mechanisms are not mutually exclusive, it remains uncertain whether their dynamics are altered in the context of disease, particularly in the context of stunted child growth.

**Figure 1:**
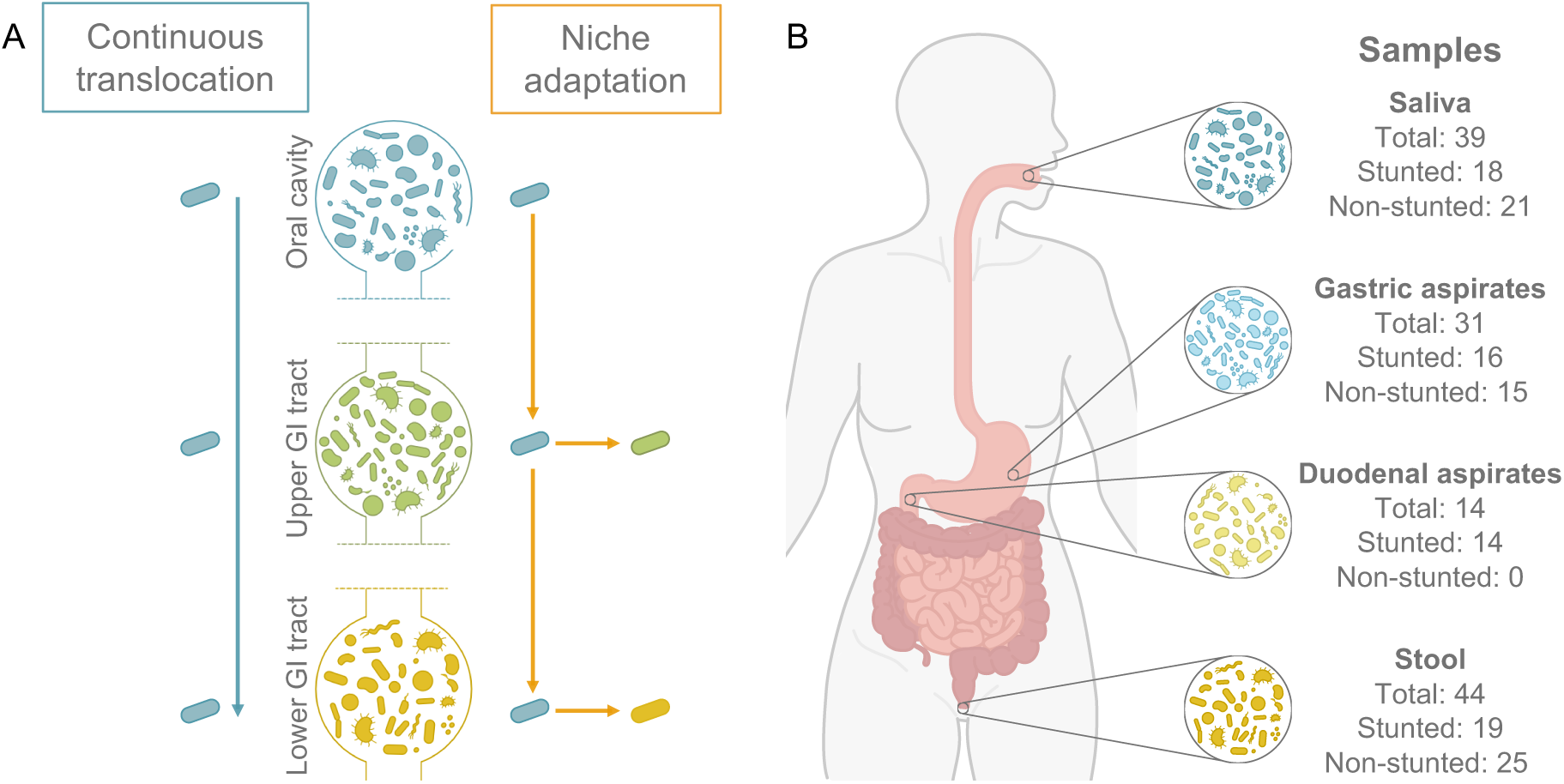
Translocation mechanisms and samples type. A. Bacterial strain translocation mechanisms from the oral cavity to the upper and lower gastrointestinal tract. Bacterial strains can continuously translocate and remain the same across the whole length of the digestive tract (continuous translocation). On the other side, bacterial strains can translocate and locally adapt to their respective niches for long-term residency (niche adaptation). B. Origin and number of samples collected.

Indeed, previous studies have highlighted oral-to-gut translocation and overrepresentation of oral bacteria in the lower GI tract in the context of various diseases, including colorectal cancer and inflammatory bowel disease^8,18,21^. Two potential mechanisms could explain the increased relative abundance of these bacteria observed in the small intestine and colon of diseased individuals. Changes in the intestinal environment, such as inflammation, as well as impaired host filtering/control and/or enhanced translocation may favor the passage and growth of invading oral bacteria, which are often facultative anaerobes, leading to an increased relative abundance of these taxa^22^. In contrast, Liao et al. observed that oral taxa reaching the lower intestinal tract have an increased relative abundance due to the overall depletion of other intestinal bacteria^22^. Distinguishing between these scenarios both in the small and large intestine will allow to develop therapeutic strategies targeting the microbiome of children suffering from stunted growth.

Here, we investigated bacterial translocation and enrichment mechanisms of oral strains along the digestive tract in both non-stunted and stunted children from Bangui, Central African Republic, using a combination of shotgun metagenomic sequencing of oral, gastric, duodenal and fecal samples and whole genome sequencing of bacterial isolates. We show that a subset of bacterial species is found both in the upper and lower GI tract and strain tracking indicates common translocation along the digestive tract. In addition, we find *Streptococcus salivarius* strain translocation events in the majority of individuals showing a potential source-sink dynamic with a continuous seeding from the oral cavity to the lower GI tract. Finally, we proposed two opposing mechanisms to explain the enrichment of oral strains observed in the small and large intestines of stunted children.

## Materials and methods

### Study cohort and sample collection

Children included in this study were recruited in the context of the TITI (*Transmission Investiguée de la Turbeculose Infantile*) project, a multicenter case-control study carried out in major urban centers in Benin, Burkina Faso, Cameroun and Central African Republic that aimed at assessing the feasibility of contact investigation and effectiveness of preventive therapy for pulmonary tuberculosis (TB) for children under 5 years of age^23^. Children recruited in the present study were exclusively from Bangui, Central African Republic and were originally enrolled in the TITI project as household contact of tuberculosis cases, without clinical signs of TB^23^. Inclusion criteria for this specific study followed the Afribiota study inclusion criteria^24^. Additionally, exclusion criteria included HIV positive results, suspicion of tuberculosis and acute malnutrition. Children were pre-selected at the TITI screening and treatment centers from among those who participated in the TITI project, given a written and oral informed consent and were invited to participate to this study. The children were then redirected to the Complexe Pédiatrique de Bangui for sample taking and for all other study procedures. The children were divided into the following groups: Moderately or severely undernourished children (height-for-age z-score ≤ −2 and weight-for-height z-score ≥ - 2) and control children with no chronic undernutrition (height-for-age z-score ≥ −2 and < +2 and weight-for-height z-score ≥ −2 and < +2). Children with acute malnutrition (weight-for-height z-score ≤ −2) or tuberculosis were excluded. No sample size estimation was performed, as the *a priori* variation in strain translocation was not known. The target size we aimed for was 40 children, divided equally between stunted and non-stunted children. Stool samples, blood samples, gastric aspirations and saliva samples were collected from all children following the same procedure as in the Afribiota project^11,24–26^. Biomarkers were measured as described for the Afribiota project^26^. In addition, duodenal aspirates were collected from stunted (height-for-age z-score < −2) children only.

### Ethical approval

Ethical clearance for the extension of the TITI (*Transmission Investiguée de la Turbeculose Infantile*) project to collect additional samples used in the present study was approved by the Scientific and Ethical Committee of the Health Science Faculty of the University of Bangui, Central African Republic, under No. 3/UB/FACSS/CSCVPER/17, 18/UB/FACSS/CSCVPR/18 and 09/UB/FACSS/CSCVPER/18. All participants received oral and written information on the study, and the caregiver provided written consent for their child to participate in the study. The project was performed following the ethical principles of the Declaration of Helsinki.

### Calprotectin and alpha1-antitrypsin measurements

Calprotectin and alpha1-antitrypsin (AAT) were used as inflammation and protein-losing enteropathy markers and were measured from 5g of stool at the coprology laboratory of the Pitié Salpêtrière Hospital according to standard accredited procedures as described previously^27^. Fecal AAT was measured using an immunonephelometric method adapted on the BN ProSpec system (Siemens) and 0.15mg/g was considered the upper limit of the normal range. Fecal calprotectin was assayed in duplicate by ELISA (Calprest, Eurospital) according to the manufacturer’s instructions. The upper limits of the normal range (no inflammation) were: < 150μg/g for children aged 24-35 months, < 100μg/g for children aged 36-59 months, and < 50μg/g for children above 59 months. Children with values outside the thresholds were considered as having intestinal inflammation or protein exudation (Supplementary Table S1).

### Sample DNA extraction

DNA was extracted from the saliva, gastric aspirates and duodenal aspirates using the Maxwell®’s RSC PureFood GMO and Authentication kit (Promega) following the manufacturer instruction with additional bead beating and enzymatic digestion steps. Briefly, samples were thawed in anaerobic environment, and 400μl of sample was centrifuged at 15,000G for 30 minutes. Pellets were resuspended in 500μl of CTAB buffer (Promega) in PowerBead Pro tubes (Qiagen), incubated at 95°C for 5 minutes and homogenized by bead beating for 10 minutes. 130μl of 2x Tissue and Cell lysis buffer (LGC Biosearch Technologies) and 5μl of 25mg/mL lysozyme from chicken egg-white (Sigma-Aldrich) were added and incubated for 30 minutes at 37°C. 40μl of proteinase K (Promega) and 20μl of RNAse (Promega) were added and the solution was incubated at 65°C for 15 minutes, then cooled down on ice for 5 minutes. The resulting solution was centrifuged at maximum speed for 5 minutes. 600μl of supernatant was then transferred into the first well of the Maxwell®’s cartridge (Promega) containing 300μl of lysis buffer (Promega). Following the Maxwell®’s run, DNA was eluted in 100μl of elution buffer (Promega).

DNA from the stool sample was extracted using QiAmp DNA Mini kit (Qiagen) following the manufacturer instruction with additional bead beating step using either 0.7-1.2mm Granat beads (BioLabProducts GmbH). Briefly, 50mg of samples were transferred to 250μl of 2% Polyvinylpolypyrolidone (PVPP) buffer (Sigma-Aldrich) directly in the bead tubes and homogenized by bead beating for 10 minutes. Samples were then frozen at −80°C for 30 minutes before being directly heated at 70°C for 2 minutes. Samples were then processed following the kit’s instructions.

Library preparation and paired-end (2×150bp) shotgun metagenomic sequencing was performed on extracted DNA by Novogene. The genomic DNA samples were fragmented into short fragments that were end-polished, A-tailed and ligated with full-length adapters for Illumina sequencing before further size selection. PCR amplification was then conducted through the AMPure XP system (Beverly), and the resulting library was assessed on the Agilent Fragment Analyzer System (Agilent) and quantified to 1.5nM through Qubit (Thermo Fisher^TM^) and qPCR. The qualified libraries were pooled and sequenced on Illumina NovaSeq 6000 (PE150) with a target depth at 40 millions reads (∼12Gbp) for the saliva, gastric aspirate, duodenal aspirate and 25 stool samples. Target depth was at 100 millions reads (∼30Gbp) for the remaining 19 stool samples.

### Metagenomic quality control, profiling and assembly

Adapter removal and quality trimming was performed using fastP (v0.23.4)^28^ and resulting reads were aligned to the human genome with Bowtie2 (v2.5.1)^29^, in –very-sensitive-local mode, for host reads removal. Taxonomic abundance profiling on the clean reads was done with MetaPhlAn4 (v4.1.1) using the ChocoPhlAn nucleotide database (SGB_202307)^30,31^.

Strain-level profiling using StrainPhlAn3^30,31^ was performed by extracting the MetaPhlAn4 marker genes and by producing the multiple sequence alignment (MSA) to build a phylogenetic tree per marker gene using PhyloPhAn^32^. Resulting phylogenetic trees were normalized by their total branch length and distance between tips was calculated using the ape package (v5.8-1)^33^ in R. Strain pairs with a distance less than 0.001 were identified as strain sharing events.

Metagenomes were assembled using metaSPAdes (v3.15.5)^34^ with -k 21,33,55,77,99,127. Assembled scaffolds longer than 1,000 nucleotides were binned into Metagenome-Assembled Genomes (MAGs) using MetaBAT2 (v2.12.1)^35^ with custom parameters --minContig 2,000, --maxEdges 500, --minCV 1 and --minClsSize 20,000. Quality control of all resulting genomes was done with CheckM (v1.1.3)^36^ and taxonomic assignment was performed using GTDB-TK (v2.4.0)^37^.

Strain-level profiling using inStrain (v1.9.0)^38^ was performed using MAGs first and subsequently, adding the assembled isolate genomes as input. Genomes were dereplicated at the strain level using dRep (v3.5.0)^39^ with parameters -pa 0.95, -sa 0.98, -nc 0.30, -comp 75, -con 10 and --S_algorithm fastANI (v1.34)^40^. Alignment was performed using Bowtie2 (v2.5.1)^29^ and genes were profiled using Prodigal (v2.6.3)^41^. A genome identity of 99.999% and coverage of 0.5 were used to identify strain sharing events.

### Selective isolations of *Streptococcus* strains

*Streptococcus* strains were isolated from saliva and stool samples from at least six stunted (height-for-age z-score < −2) individuals and six non-stunted (height-for-age z-score > −2) individuals with sufficient amount of material available. Individuals with a gastric aspirate pH above 5 or duodenal aspirates pH under 6 were excluded.

Saliva, gastric aspirates and duodenal aspirates samples were thawed in anaerobic environment and 10μl was diluted in 90μl of 1X phosphate-buffered saline (PBS, Sigma-Aldrich) solution. Stool samples were partially thawed in anaerobic environment and 0.1g of sample was diluted in 1ml of 1X PBS. Sample were thoroughly resuspended and serially diluted from 1:10 to 1:1000. Dilutions were plated on either Schaedler Anaerobe agar (Thermo Fisher^TM^), Brain Hearth Infusion agar (BHI, Thermo Fisher^TM^), BHI supplemented with 1g/L of inulin from chicory root (Sigma-Aldrich), BHI supplemented with 5g/L of yeast extract (Thermo Fisher^TM^) and 1g/L of cysteine (Sigma-Aldrich), Columbia Blood agar (CBA, Thermo Fisher^TM^) with 5% defibrinated sheep blood (Thermo Fisher^TM^), Man-Rogosa-Sharpe agar (MRS, Thermo Fisher^TM^), modified Gifu Anaerobic Medium agar (mGAM, HyServe GmbH & Co. KG) or Selective Strep agar base (Hardy Diagnostics) in aerobic and anaerobic environments. Following a 24 h incubation, colonies were re-streaked to purity on CBA plates.

Isolates were identified using either the MALDI-TOF Biotyper (Bruker-Daltonics) or by full length 16S rRNA gene amplicon sequencing. Single colonies of isolates to be identified by MALDI-TOF were fixed on two MALDI target spots using 1μl of formic acid (Sigma-Aldrich) and 1μl α-cyano-4-hydroxycinnamic acid (HCCA, Bruker Daltonics). Isolates were identified using the MALDI Biotyper reference library (Bruker-Daltonics) and the open-source CLOSTRI-TOF database^42^.

Colonies were incubated with 200μl of 5% Chelex® resin (Bio-Rad) at 95°C for 10 minutes. The bacterial 16S rRNA gene was then directly amplified from by PCR using the forward primer 27f (5′-AGAGTTTGATCCTGGCTCAG-3′) (100μM), reverse primer 1429r (5′-TACGGYTACCTTGTTACGACTT-3′) (100μM) and the Firepol PCR Mastermix 5x (Solis Biodyne). Cleaning of the PCR product was performed using the Wizard® SV Gel and PCR Clean-Up System kit (Promega) following the manufacturer instructions. Extracted DNA was sequenced by Sanger sequencing at MicroSynth AG. Forward and reverse reads were aligned using FLASH (v1.2.11)^43^ and aligned sequences were identified using BLASTn (v2.15.0+)^44^ with the 16S rRNA database downloaded on the 15.03.2024.

### Isolates sequencing and assembly

Isolates were plated overnight on BHI agar plates in aerobic condition; single colonies were then transferred into 6mL BHI liquid broth (Thermo Fisher^TM^) and incubated overnight in aerobic conditions. DNA was extracted using the Wizard® Genomic DNA purification kit (Promega) with an additional enzyme digestion step. Briefly, cell pellets were resuspended in 480μl of 50μM Ethylenediaminetetraacetic acid (EDTA, Invitrogen) and 120μl of 25mg/mL lysozyme from chicken egg-white (Sigma-Aldrich) and incubated at 37°C for 30 minutes. Samples were then processed following the manufacturer instructions.

Out of the 100 sequenced isolates, 25 were sequenced using short-reads and 75 were sequenced using long-reads. Short read DNA sequencing was performed by Novogene using Illumina NovaSeq X Plus Series (PE150). Sequencing reads were cleaned using fastP (v0.23.4)^28^ and assembled using SPAdes (v3.15.5)^45^ with -k 21,33,55,77,99,127. Long reads were sequenced using Pacific Biosciences (PacBio) Revio platform at the Lausanne Genomic Technologies Facility (GTF). HiFi reads were assembled using Flye (v2.9.3)^46^. Assembled genomes quality was assessed using CheckM (v1.1.3)^37^ and taxonomic assignment was performed using GTDB-TK (v2.4.0)^37^. Assembled genomes identified as *Streptococcus salivarius* or *Streptococcus parasanguinis* with a completeness above 95% and less than 5% contamination were retained for further analysis.

Information, such as the sample of origin, growth conditions, sequencing platform and quality, about the sequenced isolates can be found in Supplementary Table S2.

### Phylogenetic tree reconstruction and Average Nucleotide Diversity

Assembled genomes were annotated using Prodigal (v2.6.3)^41^. OrthoFinder (v2.5.5)^47^ was used to infer orthogroups and to do multiple sequence alignment based on single copy orthologs using the -M msa mode. A species phylogenetic tree was constructed using IQ-TREE2^48^ (substitution model: Q.plant+F+I+G4) with 1,000 bootstrap replicates (-bb 1000) for branch support. Four *Streptococcus parasanguinis* genomes were used as an outgroup to root the tree. Three non-focal isolate genomes of *Streptococcus salivarius* were included during orthogroup inference and tree estimation, then pruned from the rooted tree prior to visualization. In R, the package ape (v5.8-1)^33^ was used for tree manipulation and ggtree (v3.14.0)^49^ for visualization. Pairwise average nucleotide identity (ANI) between all strains was calculated using fastANI (v1.1)^40^ and an ANI > 99.9% was used as threshold to identify identical strains^50^.

### Bacterial species categories

Bacterial species were categorized based on their prevalence across samples. To categorize the species in the oral and fecal samples, only the 39 individuals with paired saliva and stool samples were considered and taxa with less than 10% prevalence and a relative abundance under 10^-4 were filtered out. Species were categorized as described in Table 1.

**Table 1:** Definitions of the species categories for comparison of samples from different compartments.

| Category | Definition |
| --- | --- |
| Uniquely oral | Present only in oral samples |
| Oral origin | $\geq 10\%$ in oral, $< 10\%$ in fecal |
| Uniquely fecal | Present only in fecal samples |
| Fecal origin | $\geq 10\%$ in fecal, $< 10\%$ in oral |
| Mostly shared | $\geq 10\%$ and $< 60\%$ in both oral and fecal |
| Highly shared | $\geq 60\%$ and $< 100\%$ in both oral and fecal |
| Completely shared | Completely shared |
| **Additional categories for oral, upper GI tract and fecal samples comparison** |  |
| Taxa shared in upper digestive tract | $\geq 10\%$ in oral and upper gastrointestinal, $< 10\%$ in fecal |
| Taxa shared in lower digestive tract | $\geq 10\%$ in upper gastrointestinal and fecal, $< 10\%$ in oral |

To categorize the species in the gastric and duodenal aspirates, only the 11 individuals with paired gastric and duodenal samples were considered and taxa with less than 10% prevalence and an abundance under 10^-4 were filtered out. Species were then categorized similarly than for the oral and fecal samples (Table 1).

To categorize the species in the oral, upper GI tract and fecal samples, only the 27 individuals with saliva, gastric and stool samples were considered. The 27 gastric aspirates and the 11 duodenal aspirates were considered as upper GI tract samples. Taxa with less than 10% prevalence and an abundance under 10^-4 were filtered out. Taxa were categorized similarly than for the oral and fecal samples with additional categories (Table 1).

### Bacterial absolute abundance

For assessing the bacterial load in the duodenal aspirates, qPCR against standard curve was used to determine CFU/mL. For each sample, triplicates of 20μl were prepared with each containing 6μl of 15-25ng/μl of extracted DNA, 7.6μl of SYBR Green PCR Master Mix (Life Technologies) and 1.4μl of forward (5’-YCGTACTCCCCAGGCGG-3’) and reverse (5’-AGGATTAGATACCCTGGTAGTCC-3’) primers at 4μM amplifying the 16S rRNA gene. A standard curve was established using genomic DNA extracted from *Streptococcus salivarius* strain AF23 at 10^4^, 10^5^ and 10^6^ CFU/mL. Cycling conditions of the qPCR were 50°C for 2 min, 95°C for 10 min, followed by 40 cycles of 95°C for 15 s and 60°C for 1 min followed by a cycle to determine the melting curve. The number of CFU/mL for each sample was extrapolated from the mean Ct value of the triplicates using the results from the standard curve and multiplying the final results by the DNA dilution factor.

To obtain an estimation of the absolute bacterial biomass in the feces we used a published method that estimates the bacterial-to-host read count ratio^51^. For each fecal sample, the natural logarithm of the number of assigned bacterial reads divided by the number of host reads removed was computed. Sample AGN001 was considered an outlier and removed from the analysis due to the presence of more host reads than bacterial reads. The relationship between the absolute bacterial biomass and predictors such as height-for-age z-score, sex, age, mode of delivery, or intestinal inflammation was assessed by linear regression analysis.

### Nucleotide diversity and popANI of *Streptococcus salivarius*

MAGs and isolate genomes were dereplicated using dRep (v3.5.0)^39^ with parameters -pa 0.90, -sa 0.95, -nc 0.30, -comp 75, -con 10 and --S_algorithm fastANI (v1.34)^40^. Alignment was performed using Bowtie2 (v2.5.1)^29^ and genes were profiled using Prodigal (v2.6.3)^41^. InStrain profile^38^ was used, and the output table were filtered to retain only the profiling of *Streptococcus salivarius* reference genome. Using option – genome, the *Streptococcus salivarius* genome was used in inStrain compare^38^ to obtain the popANI within and between samples.

### Statistical analyses

Statistical analysis and visualization were conducted in R (v4.2.2)^52^ using packages gtsummary (v2.2.0)^53^, ggplot2 (v3.5.1)^54^, phyloseq (v1.50.0)^55^, vegan (V.2.6-10)^56^, MicEco (v0.9.19)^57^, ComplexHeatmap (v2.22.0)^58^, microbiomeMarker (v1.12.2)^59^, ANCOMBC (v2.8.1)^60^, geiger (v.)^61^ and ape (v5.8-1)^33^.

Alpha diversity was assessed using the estimate_diversity function from the phyloseq^55^ package using the Observed and Shannon indexes, and groups were compared using the Wilcoxon rank-sum test with Benjamini-Hochberg (BH) correction for multiple testing. Beta diversity was assessed using Principal Coordinate Analysis (PCoA) based on Bray-Curtis distance. Community composition difference across conditions was assessed with PERMANOVA (Permutational Multivariate Analysis of Variance) using the adonis2 function from the vegan^56^ package. The envfit function from the vegan^56^ package was used to assess which environmental variables explained the clustering of the samples by fitting environmental vectors such as body site, sex, mode of delivery, intestinal inflammation or nutritional status, on the ordination space. Differential abundance analysis between categorical variables was evaluated using the run_ancombc and run_lefse functions from the microbiomeMarker^59^ package with default parameters and BH correction for multiple testing. The function ancombc with default parameters from the ANCOMBC^60^ package was used to assess for association between bacterial species and the height-for-age z-score (HAZ) and p-values were adjusted for multiple testing with the BH method.

Transition rates between compartments and between individuals were inferred using maximum likelihood macro-evolutionary Markov models for discrete character evolution^62,63^. Models were fitted using the observed phylogeny and the distribution of bacterial features (either individual of origin or compartment of origin) at the tip of the phylogeny. Model for compartment and model for individuals IDs were independently fitted using the geiger^61^ R package with the function fitDiscrete under an equal-rates (ER) model. The phylogeny included solely *Streptococcus salivarius* isolates. Initial fit of the model showed extremely low likelihood for the compartment trait (log Lik −10e-200), likely due to the existence of extremely short branch lengths of strains from different compartments. In this case, transition rates between compartment will be extremely high as transitions occur on extremely short branches, making estimation of transition rates uncertain. To mitigate this problem, the tree was transformed into an ultrametric tree using the chronos function from the ape^33^ package under a relaxed clock model (Supplementary Figure S1). Transition rates between states were estimated from an ER model as above. The ratio of estimated compartment-to-individual transition rates was then obtained to quantify relative transition rates. Because our ultrametric tree shows less extreme small branch lengths than the untransformed tree (Supplementary Figure S1), our ratio estimate will be highly conservative, i.e. the true ratio is probably higher than the one estimated with the ultrametric tree. If the observed ratio is still higher than expected by chance, this means that our conclusion will also be highly conservative. To assess whether the observed ratio could be explained by chance only, trait labels were randomized across tips 499 times and we refitted the ER model for both traits on each randomized dataset. For each replicate, the ratio of compartment-to-individual transition rates was recorded. The null distribution of randomized ratios was compared to the observed ratio, and a p-value was obtained as the fraction of random replicates exceeding the observed value.

### Code and data availability

Scripts used in the data processing and analysis are available via GitHub (https://github.com/VonaeschLabUNIL/Afribiota/tree/main/Satellite).

Original sequencing data is protected due to ethical constraints and should be directly requested to the Afribiota consortium through the lead author Pascale Vonaesch. Dereplicated (>98% ANI) Metagenome Assembled Genomes (MAGs) generated from this study and sequenced genomes from isolates have been deposited in the European Nucleotide Archives (ENA) at EMBL-EBI under accession number PRJEB102291.

## Results

### Microbiome diversity and composition differs along the digestive tract

Out of the 44 included participants, 25 followed a normal growth (height-for-age z-score >= −2) and 19 suffered from stunted growth (height-for-age z-score < −2) (Table 2). Collected samples included saliva (*n*=39), gastric aspirates (*n*=31) and stool (*n*=44) for all recruited children, as well as duodenal aspirates (*n*=14) from stunted individuals (Table 2). In total, 128 samples were subjected to deep metagenomic sequencing (Figure 1B). As expected, body compartment was a major driver of species richness, with the stool samples showing significantly higher richness than the other body sites (saliva *p*-value 2.08e-6, r = 0.53, gastric *p*-value 6.30e-11, r = 0.79, duodenum *p*-value 5.15e-6, r = 0.61). Saliva samples showed a significantly higher species richness than the gastric (*p*-value 8.76e-8, r = 0.66) and the duodenal samples (*p*-value 6.00e-3, r = 0.39). Saliva and duodenal samples showed higher evenness measured by the InvSimpson index than the gastric samples (saliva *p*-value 1.50e-2, r = 0.33, duodenum *p*-value 3.80e-2, r = 0.33) and fecal samples (saliva *p*-value 1.00e-3, r = 0.40, duodenum *p*-value 1.50e-2, r = 0.35) (Figure 2A, Supplementary Table S3). Sample diversity measured by the Shannon index showed no significant differences between each body site (Figure 2A, Supplementary Table S3).

**Figure 2:**
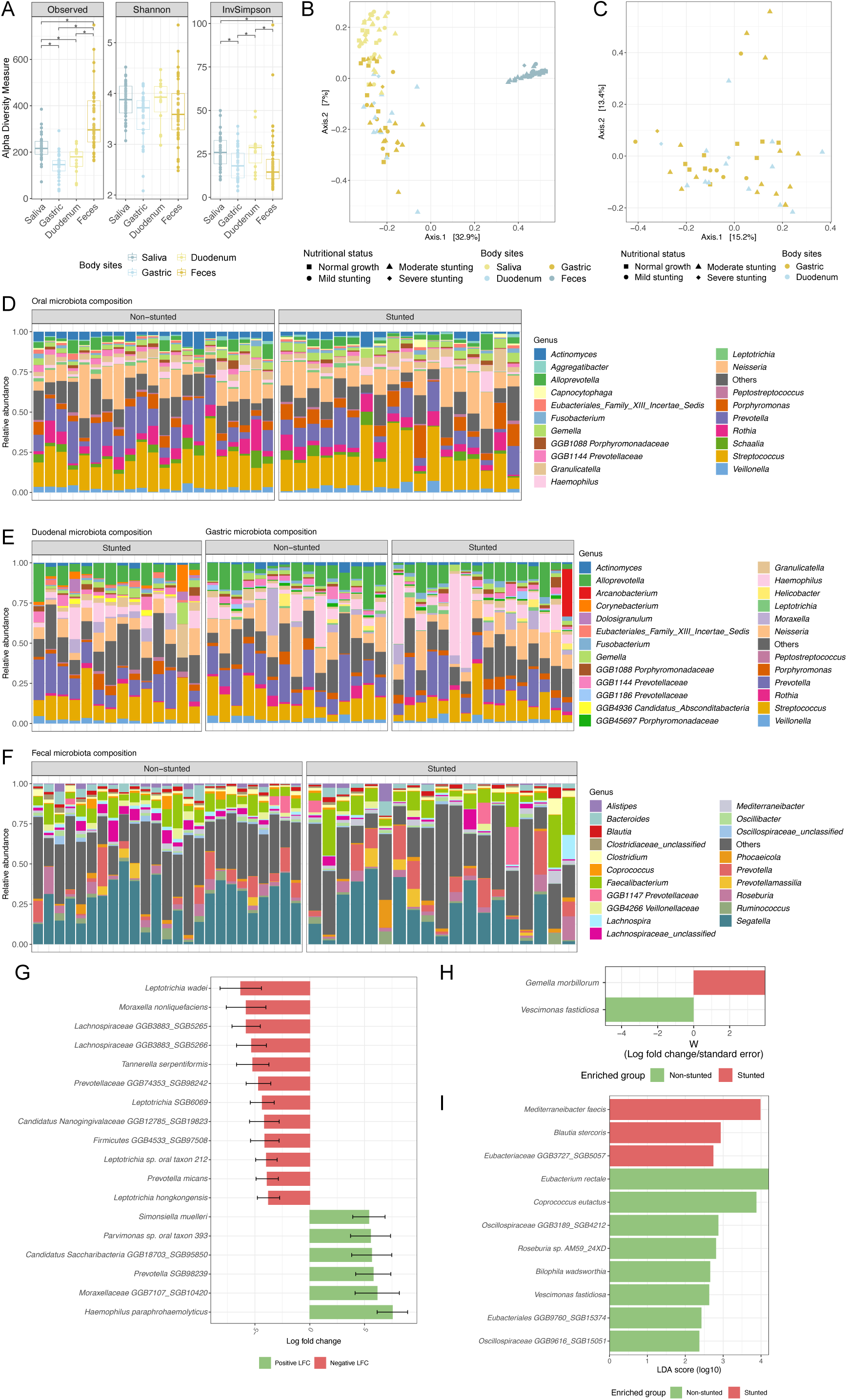
Microbiota diversity and composition differs between body sites and nutritional status. A. Alpha-diversity metrics (Observed and Shannon indexes) comparing the four body sites. Pairwise comparisons using Wilcoxon rank sum test with Benjamini-Hochberg correction for multiple comparisons. B. Principal Coordinates Analysis (PCoA) based on Bray-Curtis distance, showing ordination of samples. C. PCoA based on Bray-Curtis distance showing ordination of the gastric and duondenal samples. D-F. Bacterial genera relative abundance in the (D) saliva, (E) gastric and duodenal, and (F) stool samples. Samples split by stunting status. Top 20 most abundant genera in each compartment, remaining genera are grouped in the “Others” category. G. Association between bacterial species and height-for-age z-score (HAZ) in the duodenal samples tested with ANCOM-BC with Benjamini-Hochberg correction for multiple testing. Black bars indiciate standard errors. H. Enrichment of bacterial species in the feces of stunted and non-stunted children tested with ANCOM-BC with Benjamini-Hochberg correction for multiple testing. W (Wald statistic): log fold change/standard error. I. Linear discriminant analysis (LDA) Effect Size (LefSe) analysis showing the enrichment of bacterial species in the feces of stunted and non-stunted children.

**Table 2:** Cohort summary statistics.

| Variable | Overall $N = 44^1$ | Non stunted $N = 25^1$ | Stunted $N = 19^1$ |
| --- | --- | --- | --- |
| Age [months] | 39 (32, 51) | 41 (31, 51) | 37 (33, 51) |
| Age group |  |  |  |
| 2-3 years old | 18 (41%) | 10 (40%) | 8 (42%) |
| 3-4 years old | 12 (27%) | 6 (24%) | 6 (32%) |
| 4-5 years old | 13 (30%) | 8 (32%) | 5 (26%) |
| 5 years old | 1 (2.3%) | 1 (4.0%) | 0 (0%) |
| Sex |  |  |  |
| Female | 27 (61%) | 16 (64%) | 11 (58%) |
| Male | 17 (39%) | 9 (36%) | 8 (42%) |
| Delivery mode |  |  |  |
| Vaginal | 42 (95%) | 24 (96%) | 18 (95%) |
| C-section | 2 (4.5%) | 1 (4.0%) | 1 (5.3%) |
| Alpha1-antitrypsin (AAT) |  |  |  |
| Exudation | 7 (16%) | 2 (8.0%) | 5 (26%) |
| No exudation | 37 (84%) | 23 (92%) | 14 (74%) |
| Calprotectin |  |  |  |
| Inflammation | 13 (30%) | 7 (28%) | 6 (32%) |
| No inflammation | 31 (70%) | 18 (72%) | 13 (68%) |

<sup>1</sup>Median (Q1, Q3); n (%)
Stunting threshold: height-for-age z-score < -2
AAT exudation threshold: > 0.15mg/g
Calprotectin inflammation thresholds: $\geq 150\mu\text{g/g}$ for children aged 24-35 months, $\geq 100\mu\text{g/g}$ for children aged 36-59 months, and $\geq 50\mu\text{g/g}$ for children above 59 months

The clustering of the samples on a PcoA plot based on Bray-Curtis’s distance showed a clear separation by body compartment (R^2^ = 0.375, *p*-value 1e-4), with exception of the gastric and duodenal samples (Figure 2B, Supplementary Table S3). These samples exhibit high compositional similarities (*p*-value 0.6933, Figure 2C). To further investigate factors associated with beta-diversity, additional covariates, such as the nutritional status, mode of delivery or intestinal inflammation were assessed. None of these variables were found to be significantly associated with microbiome composition (Supplementary Table S3).

Saliva samples yielded an average of 25.3 million high quality reads and were largely dominated by *Streptococcus* species (20.7% mean relative abundance), followed by *Neisseria* spp. (12.6%) and *Prevotella* spp. (12.1%) (Figure 2D, Supplementary Table S3). The most dominant *Streptococcus* species were *S. mitis*, *S. salivarius* and *S. parasanguinis* with a mean relative abundance of 5%, 4.2% and 3.8%, respectively (Supplementary Table S3). The most abundant species were *Neisseria subflava* and *Rothia mucilaginosa* representing 6.6% and 5.2% relative abundance in the saliva samples (Supplementary Table S3).

Gastric and duodenal aspirates resulted in an average of 4.9 and 8.8 million high quality reads, respectively (Supplementary Table S3). These two body sites were largely dominated by *Prevotella* (13.6% in the gastric samples and 12.7% in the duodenal samples), *Streptococcus* (12.9% and 15.6%), *Alloprevotella* (12.6 and 9%), *Neisseria* (11.2% and 8.3%) and *Haemophilus* (9.3% and 6.6%) (Figure 2E, Supplementary Table S3).

The stool samples were sequenced at two targeted sequencing depths. The first subgroup of 19 samples generated an average of 110 million high quality reads while the second subgroup of 25 samples resulted to an average of 43.9 million high quality reads. No significant differences in richness or diversity were detected between these two groups of samples (Supplementary Table S3). Stool samples were largely dominated by *Segatella* species, that represented a mean relative abundance of 24%, notably *Segatella copri* with a 10.5% mean relative abundance (Supplementary Table S3). Other dominant taxa include *Faecalibacterium* (8.9%), *Prevotella* (6.5%) and *Roseburia* (3.4%). *Streptococcus* represented only 0.6% of the mean relative abundance in the stool samples (Figure 2F, Supplementary Table S3).

To identify taxa associated with stunting severity in the duodenum, the height-for-age z-score (HAZ) as a continuous variable was used as a covariate in the differential abundance analysis based on read count using the Analysis of Compositions of Microbiomes with Bias Correction (ANCOM-BC) at the species level (Supplementary Table S3). This approach showed a significant negative association between HAZ and 12 species, including multiple members of the *Leptotrichia* genus, *Moraxella nonliquefaciens*, *Tannerella serpentiformis*, and *Prevotella micans* (Figure 2G). Conversevly, six species were postivily associated with HAZ in the duodenum (Figure 2G). In the feces, no significant associations were found between HAZ and bacterial species abundance (Supplementary Table S3). Nevertheless, using the binary stunting classification, differential abundance analysis using ANCOM-BC showed an enrichment of *Gemella morbillorum* in stunted children (HAZ <= −2) and an enrichment of *Vescimonas fastidiosa* in non-stunted children (HAZ > −2) (Figure 2H). Furthermore, using Linear Discriminant Analysis (LDA) Effect Size (LefSe) for differential abundance analysis based on relative abundance, stunted children showed a depletion of butyrate-producing bacteria, such as *Eubacterium rectale* and *Coprococcus eutactus* in their feces (Figure 2I).

### A subset of species is found along the whole gastrointestinal tract

To evaluate the compositional differences between the microbiome of the different body compartments and to detect for microbial translocation along the digestive tract, bacterial species were classified into categories based on their prevalence across subjects (See Materials and Methods, Figure 3A).

**Figure 3:**
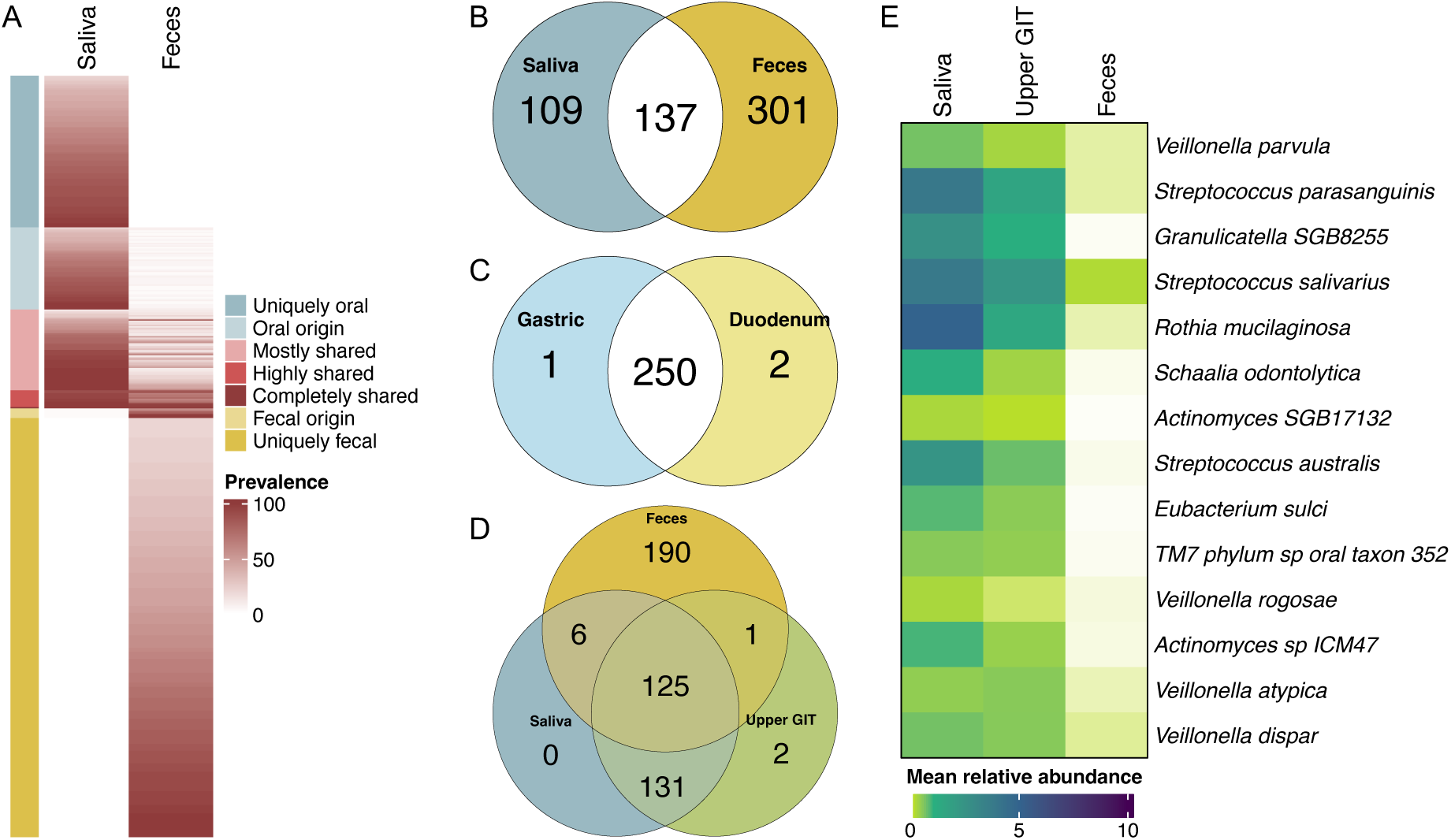
A subset of bacterial species are shared across all body compartments. A. Prevalence of all species in the saliva and fecal samples. Species were categorized based on their prevalence in each body site. B. Number of bacterial species unique to the saliva or fecal samples, and number of bacterial species found at least once in both body sites. C. Number of bacterial species unique to the gastric or duodenal aspirates, and number of bacterial species found at least once in both body sites. D. Number of bacterial species unique to the saliva, upper gastro-intestinal tract (GIT) and fecal samples, and number of species shared between these different body sites. E. Mean relative abundance of the 14 highly shared species (>60% prevalence) between the saliva, upper GIT, and fecal samples.

In the 39 paired saliva and stool samples, a total of 547 species were detected (prevalence >= 10% and abundance > 10e-4), of which 109 were uniquely found in the oral cavity, representing roughly 22% of the relative abundance, while 301 were uniquely found in the feces and accounting for more than 83% of the relative abundance (Figure 3B). In addition, 59 species were classified as being of oral origin (observed in <10% of fecal samples, ∼20% of the oral relative abundance) and 7 as being of fecal origin (observed in <10% of saliva samples, ∼3% of the fecal relative abundance) (Figure 3A). Predominantly oral taxa included several *Prevotella* species such as *P. melanonigenica*, *P. nanceiensis*, *P. pallens* and *P. histicola*, as well as *Porphyromonas bobii, Neisseria sicca, Haemophilus haemolyticus* and an *Alloprevotella* species all found in more than 80% of the saliva samples and with a relative abundance above 1% (Supplementary Table S3). Amongst the 301 species classified as being predominantly fecal, the most abundant (>1%) and prevalent (>80%) ones were *Segatella copri*, *Faecalibacterium prausnitzii*, *Roseburia inulinivorans* and *Mediterraneibacter* [*Ruminococcus*] *faecis* (Supplementary Table S3).

Additionally, 71 bacterial species were classified as being shared between the two compartments based on their prevalence across the paired samples, accounting for more than 55% of the oral relative abundance and less then 1% of the fecal (Figure 3A). Of note, no *Prevotella* or *Segatella* species were found to be shared between the oral cavity and the feces, even though these bacterial genera represented a high relative abundance in both body sites (Supplementary Table S3).

Most taxa were found to be shared in the 11 paired gastric and duodenal aspirates (231/253 shared taxa, prevalence > 10% in both body sites) and accounted for more than 93% of the relative abundance at both body sites (Figure 3C). Amongst these species, *Helicobacter pylori*, found in 12 out of 31 gastric aspirates (relative abundance 1.25%) and *Moraxella nonliquefaciens* were the only taxa never found in the saliva or stool of the 27 children who provided at least three biological samples (Supplementary Table S3).

Out of the 455 detected species (prevalence >= 10% and abundance > 10e-4) in the saliva, upper GI and stool samples, 68 were found to be shared, accounting for roughly 54%, 27% and 0.6% relative abundance, respectively (Figure 3D, Supplementary Table S3). Members of the *Veillonella*, *Streptococcus*, *Rothia*, *Actinomyces*, *Eubacterium*, and *TM7* genera were amongst the 14 highly shared species (> 60% prevalence) across all body sites (Figure 3E). Importantly, among these highly shared species, *S. salivarius* was found to be the only bacterial species to be detected in all oral (relative abundance 4.19%) and fecal (relative abundance 0.38%) samples, as well as all the other samples (relative abundance in gastric aspirates: 2.29%, relative abundance in duodenal aspirates: 1.18%), except for one gastric aspirate (Figure 3E).

### Metagenomic resolved methods identify translocation from the upper and lower digestive tract

After observing that several species were found all along the gastrointestinal tract, StrainPhlAn3^30^ was used to investigate intra-individual strain-sharing events in between the different body compartments. Based on the multiple sequence alignment (MSA) of the dominant consensus sequence from species-specific marker genes, StrainPhlAn3 identified 38 strain-sharing events involving 23 distinct species across 19 individuals (Supplementary Table S4). Most of the events (n=35) occurred in the upper digestive tract and 23 of the 38 events involved members of the *Prevotellaceae* families, including species of *Prevotella* and *Alloprevotella* (Figure 4A). Evidence of strain translocation from the oral cavity to the stool was limited to two individuals, one stunted and one non-stunted, both of whom shared *S. salivarius* between these two compartments (Figure 4A-B).

**Figure 4:**
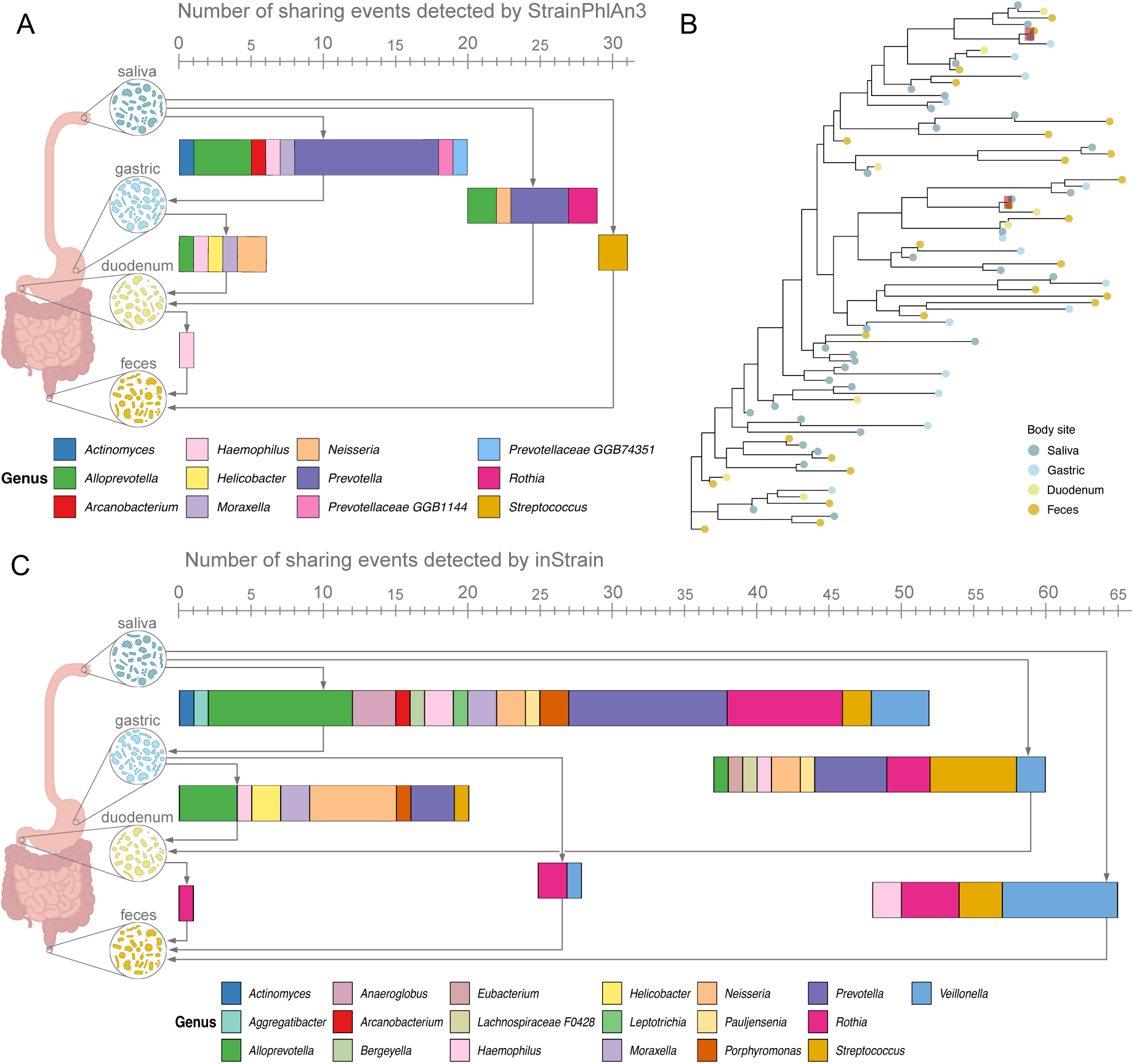
StrainPhlAn3 and inStrain detected translocations between the upper and lower digestive tract within the same individual. A. Number of sharing events detected by StrainPhlAn3, separated by paired body sites. The length of the barplots matches the number of detected sharing events. Bacterial strains are grouped into bacterial genera. B. Normalised phylogenetic tree of *Streptococcus salivarius* reconstructed from multiple sequence alignment of species-specific marker genes across shotgun metagenomic sequenced from all samples. Red-highlighted tips show the pair of samples in which a sharing event was identified (normalized phylogenetic distance < 0.001). C. Number of sharing events detected by inStrain, separated by paired body sites. The length of the barplots match the number of detected sharing events. Bacterial strains are grouped into bacterial genera.

As StrainPhlAn3 relies solely on marker genes and phylogenetic distances of consensus sequences to infer strain-sharing, inStrain^38^ was used to further assess for strain translocation based on recovered MAGs. A total of 3,221 MAGs (>=75% completeness and <10% contamination) were reconstructed and dereplicated with dRep^39^ resulting in 2,147 clusters at 98% ANI. These clusters were used as reference database for inStrain comparisons (Supplementary Table S4).

In total, 97 strain-sharing events across 27 individuals were detected by inStrain, including species of *Prevotella*, *Alloprevotella*, *Rothia*, *Streptococcus* and *Neisseria*. Of note, inStrain detected a majority of the sharing events between the saliva and the upper GI tract (Figure 4C). Six children had the same strain of either *Rothia* or *Veillonella* identified in their oral cavity, upper GI tract and feces (Supplementary Table S4). Between the oral cavity and the stool, inStrain detected 17 strains shared in 11 individuals (Figure 4C), including two *S. salivarius* strains observed in one non-stunted individual. Additionally, inStrain identified *Veillonella*, *Rothia*, and *Haemophilus* strains translocating from the oral cavity to the feces (Figure 3C).

Overall, metagenomic based approaches identified strain sharing between the oral cavity and the upper GI tract and a limited number of strain-sharing events per individuals between the upper and lower digestive tract. Evidence of *S. salivarius* translocation was observed in two children, with one case consistently detected by both StrainPhlAn3 and inStrain.

### Strain isolation enhances the recovery of *Streptococcus salivarius* genomes for assessing strain translocation events and mechanisms

To comprehensively assess for the translocation of bacterial strains along the digestive tract and to complement the metagenomic data, isolations of *Streptococcus* species targeting *S. salivarius* were performed. A total of 474 *Streptococcus* isolates were recovered from 53 samples from 23 individuals that met the eligibility criteria (see methods). From these, 87 *S. salivarius* isolates had their genome sequenced (24 by long-read sequencing and 63 by short-read sequencing) to near completeness (completeness >= 95% and contamination < 5%) and were included for analysis to obtain up to three strains isolated from at least the saliva and the feces from each individual from a minimum of five stunted and five non-stunted children.

Overall, for 11 individuals, 6 stunted and 5 non-stunted, at least one strain of *S. salivarius* was isolated from both saliva and stool. Genomes sequenced by long-read sequencing had an average genome size of 2.3Mb with an average of 1.63 fragments and an average of 149.5-fold genome coverage. *S. salivarius* genomes sequenced by short-read sequencing had an average genome size of 2.2Mb with an average of 25.08 scaffolds and an average 838-fold genome coverage (Supplementary Table S2).

In addition to the isolated genomes, 13 MAGs with completeness above or equal to 75% and contamination under 10% were reconstructed from 13 samples coming from 11 children (Supplementary Table S4). Notably, eight of these MAGs originated from samples in which no isolates were obtained. The remaining five MAGs, reconstructed from samples with isolates, showed high similarity (>98% ANI) to at least one isolates from the same sample (Figure 5A, Supplementary Table S4).

**Figure 5:**
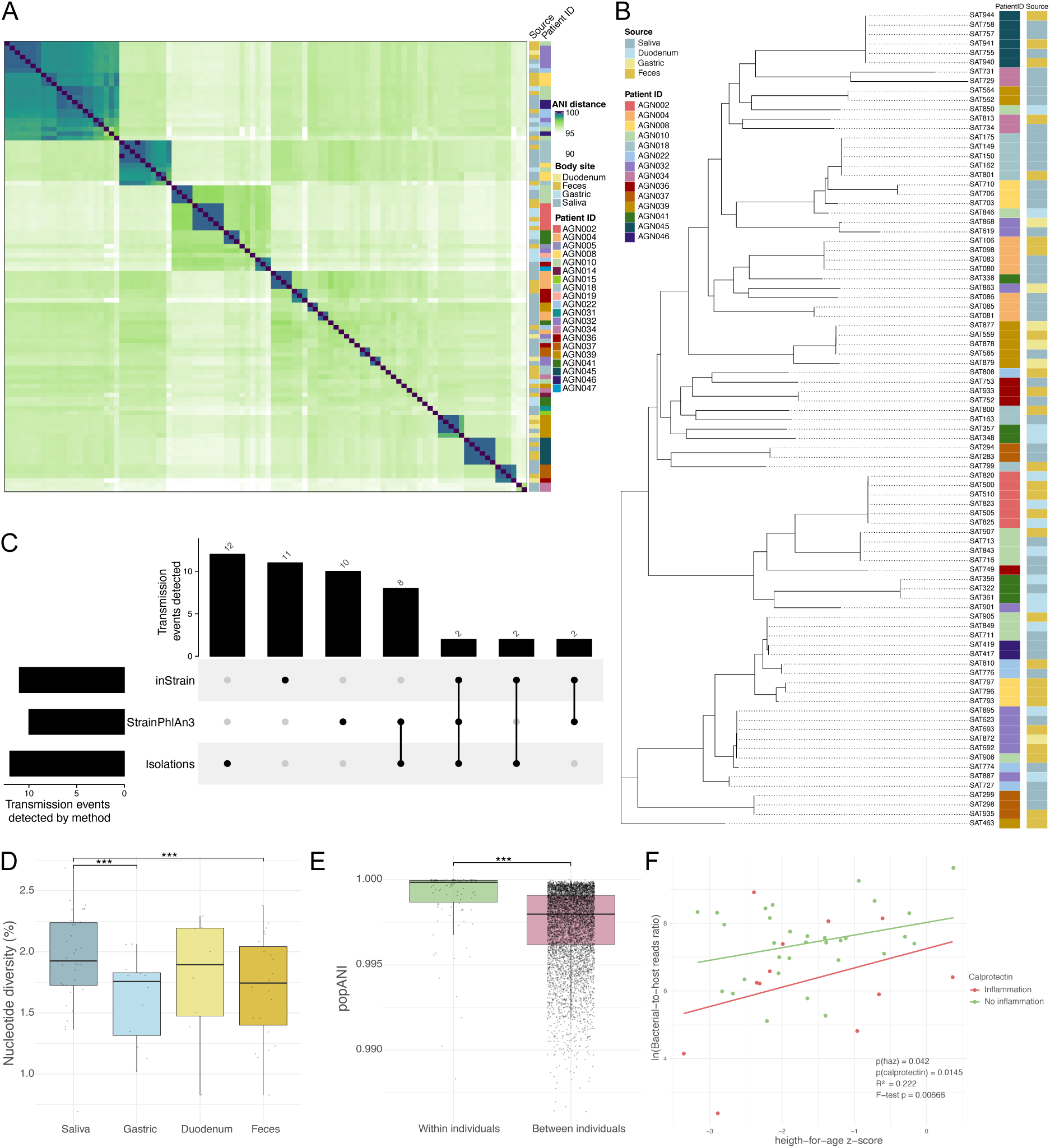
Translocation of *Streptococcus salivarius* strains is common along the digestive tract and follows a source to sink dynamic. A. Heatmap of average nucleotide identity (ANI) of the 87 *Streptococcus salivarius* isolates from the present study and the 13 reconstructed metagenome-assembled genomes (MAGs). Darker shades show higher pairwise ANI distance between strains B. Phylogeny of the 87 isolated *Streptococcus salivarius* genomes. C. UpSet plot showing the number of translocation events detected for each method and their overlap. D. Percentage of nucleotide diversity of *Streptococcus salivarius* reference genome SAT081 between the different body sites obtained using inStrain profiles. E. Population average nucleotide identity (popANI) of *Streptococcus salivarius* reference genome SAT081 within and between individuals obtained using inStrain compare. F. Association between the total biomass in the feces measured by the natural logarithm of the bacterial-to-host reads ratio and the height-for-age z-score. The effect of inflammation, measured by calprotectin levels, is depicted based on the following inflammation thresholds: >= 150μg/g for children aged 24-35 months, >= 100μg/g for children aged 36-59 months, and >= 50μg/g for children above 59 months. Statistical tests were performed with Wilcoxon rank sum test without correction for multiple testing. ***: *p*-value<0.05

Based on the relative abundance of *S. salivarius* in the samples and the sequencing depth employed, cultivation-based isolation was more sensitive at recovering and identifying multiple high-quality genomes compared to shotgun metagenomic sequencing, yet both techniques are complementary to assess for strain translocation.

### *Streptococcus salivarius* isolates show consistent translocation from the oral cavity to the lower gastrointestinal tract

Based on the isolated genomes, highly similar strains (ANI > 99.9%) from the same lineage were found in multiple compartments within the same individual, indicating 10 potential translocation events between the oral cavity and the stool in 9 children (Figure 5A-B, Supplementary Table S4). For three of these children, identical strains were also found in the gastric aspirates and for one child the same strain was found in all body compartments (Figure 5A-B).

Combining *S. salivarius* isolate sequences with metagenomic sequencing data, new translocations were detected by inStrain and StrainPhlAn3 (Figure 5C). The isolation method detected the most translocation events of *S. salivarius* strains (*n*=12) followed by inStrain (*n*=11) and finally StrainPhlAn3 (*n*=10) (Figure 5C). Overall, evidence of *S. salivarius* strains translocation from the oral cavity to the upper GI tract was identified in seven individuals and, from the oral cavity to the feces, similar strains were found in 10 individuals (Supplementary Table S4). In two individuals (one stunted and one non-stunted child), for whom isolates from multiple body compartments were available, we detected no translocation events. Additionally, for three individuals, translocation of two different strains was observed.

Integrating both metagenomic strain-tracking and isolated genomes thus allowed to consistently identify *S. salivarius* as a translocating strain in both healthy and stunted children.

### *Streptococcus salivarius* translocation follows a source-sink dynamic

Two alternative hypotheses were evaluated for the origins of *S. salivarius* populations in the lower GI tract. These populations may be (1) durably established and have specifically adapted to this compartment or (2) constantly seeded from population established in the oral cavity (source-sink dynamic within each individual). These two scenarios are expected to produce contrasting phylogenetic patterns. In the first case, lower GI tract populations are expected to form large monophyletic clades composed of strains from multiple individuals and the transition rate between oral cavity and lower GI tract compartments should be much lower compared to transition between individuals. In the second case, lower GI tract strains are expected to be closely related to strains established in the oral cavity from which they originated, and the transition rate between oral cavity and lower GI tract compartments should be much higher compared to the transition rate between individuals.

Mapping both the individuals IDs and the compartments onto the phylogeny showed that lower GI tract strains are often closely related to strains from the oral cavity, and strains isolated from a given individual tend to form large clades, favoring the second scenario (Figure 5B). To quantitatively test this hypothesis, the transition rates between compartments and between individuals were inferred using Markov models that were fitted to the observed phylogeny and trait distribution at the tip. The transition rate between compartments was 17-fold higher than between individuals, indicating that upper to lower GI tract transition happens far more often than between individuals. Randomizations of the traits and computation of the ratio to control for unforeseen biases showed a median of 1 and *p*-value of 0.019, confirming that the observed ratio is unlikely due to chance alone.

Additionally, the nucleotide diversity was evaluated across samples using inStrain profile on a *S. salivarius* reference genome (SAT081). Diversity varied by body site, with the highest levels observed in saliva (gastric *p*-value=1.25e-4, feces *p*-value=8.70e-4), followed by duodenal samples (not significant) suggesting greater strain-level heterogeneity in the oral cavity compared to the lower gastrointestinal tract (Figure 5D). To further assess population structure, population average nucleotide diversity (popANI) values were compared within and between individuals across all body sites (Figure 5E). *S. salivarius* strains from the same individual showed higher popANI than those from different individuals (*p*-value=2.2e-16), indicating that *S. salivarius* populations are more closely related within hosts than across hosts.

Together, these results were consistent with a source-sink dynamic model in which *S. salivarius* populations in the lower gastrointestinal tract are continuously seeded from the oral cavity rather than durably established. The oral cavity thus appears to act as a reservoir of strains diversity and maintains individual-specific populations along the digestive tract.

### Small and large intestines show different enrichment mechanisms of oral taxa

Lastly, potential enrichment mechanisms of taxa of oral origin in the upper and lower GI tract were examined by quantifying the total microbial load in the duodenum and feces. In the duodenal aspirates, the absolute microbial loads were quantified by quantitative polymerase chain reaction (qPCR) to obtain the total copy number of 16S rRNA genes as a proxy for colony forming unit (CFU) and to assess for the presence of SIOBO. All measured samples from stunted children had a microbial load above 10^5^ CFU/mL suggesting bacterial overgrowth (Supplementary table S4). The presence of SIOBO in the duodenum of stunted children suggests that oral bacteria enrichment may be due to a change in the intestinal environment, favoring the growth of oral bacteria in this compartment.

The total biomass in the feces was estimated using the bacterial-to-host ratio. The analyses were corrected for inflammation, as inflammation-induced host cell shedding might change these values (Supplementary table S4). Indeed, children with lower gastrointestinal tract inflammation (mean host reads removed= 695.758) had significantly more host reads removed compared to children without inflammation (mean host reads removed= 52.23, mean difference=643.519 reads, *p*-value=0.036), consistent with a previous report on 18S rRNA sequencing in a cohort of almost 1000 children from a similar context^64^. Therefore, lower gastrointestinal tract inflammation was included as covariate to evaluate the link between total biomass and stunting. Fecal bacterial biomass was positively associated with HAZ (*p*-value=0.042, R^2^=0.222) showing a reduction of overall bacterial biomass linked with stunting severity (Figure 5E). This observation suggests that, in the colon, the enrichment of oral bacteria may be due to overall depletion of other intestinal bacteria, such as butyrate-producers (Figure 2G-N), contrasting with the mechanism observed in the small intestine.

## Discussion

Ectopic colonization by bacterial strains from the oral cavity to the lower GI tract is increasingly recognized as a major contributor to the development of GI diseases^8,21^. Yet, the magnitude and mechanisms of oral-to-gut translocation remains largely uncharacterized. Ectopic oral colonization manifesting as SIOBO has been described in stunted children across many research studies and geographic regions^10–12,14,65–67^. Similarly, an overrepresentation of oral taxa has been observed in the lower GI tract of undernourished children in these regions^10,68^. Nevertheless, to date, the underlying mechanisms leading to these phenomena remain elusive. Here, we demonstrate that oral strains are found along the entire length of the digestive tract, forming a continuous population, both in healthy and stunted children. Using *S. salivarius* as model species due to its presence in all body compartments, we show that the oral cavity acts as a reservoir of strains continuously seeding the lower GI tract. Finally, we suggest two opposing mechanisms to explain the enrichment of oral bacteria in the upper and lower GI tract with overgrowth in the small intestinal tract and overall biomass reduction in the colon.

In recent years, the translocation and colonization of oral bacteria in the lower GI tract has been debated and mostly relied on 16S rRNA gene amplicon sequencing^7,69–71^. Harnessing deep shotgun metagenomic sequencing, we show that a subset of bacterial species, including members of the *Actinomyces*, *Rothia*, *Streptococcus* and *Veillonella* genera, are found along the entire length of the GI tract. These findings are in line with the results from Schmidt et al. who demonstrate frequent oral-fecal translocation of these core oral taxa in 470 individuals from five countries using shotgun metagenomic sequencing^7^. Using strain-level analysis, we further demonstrate that the same bacterial strains are detected along the GI tract although to a limited extent. Due to the low relative abundance of oral strains in the feces, the sequencing depth employed and the presence of multiple co-existing strains which may not allow proper coverage of the strains of interest, metagenomic-based methods such as StrainPhlAn3^30^ and inStrain^38^ may overlook translocation events^72^. Therefore, to complement the metagenomic data and increase the sensitivity of detecting translocation events, we isolated and sequenced strains of *S. salivarius*.

*S. salivarius* is one of the most abundant oral bacteria and was frequently shared in our study between the oral cavity and the lower GI tract. In the context of childhood undernutrition, *S. salivarius* was previously shown to be associated with a decreased lipid absorption both in cell culture and mice and was amongst the species overrepresented in the feces of stunted children from Central African Republic^10,11^. In our study in which the sample sized was restricted, differential abundance analysis did not reveal an enrichment of *Streptococcus* species in the small intestine or colon of the stunted children included. However, *S. salivarius* was found to be the sole bacterial species present in all but one of the samples. Based on ANI and phylogeny, we demonstrated that similar strains are isolated from different body compartments within the same individuals, suggesting a frequent translocation in both stunted and healthy children.

We further evaluated two possible scenarios to determine the origin of *S. salivarius* strains in the lower GI tract. Indeed, Gough et al.^19^ showed that *S. salivarius* strains from the stool of Zimbabwean children were distinct from common genomes of reference obtained predominantly from the oral cavity of individuals from high-income countries, suggesting that *S. salivarius* populations durably established and adapted to the intestinal environment. In this case, each compartment of the digestive tract would host a genetically distinct bacterial population as also suggested by Rashidi et al., who compared marker genes between 500 paired oral and fecal samples from four healthy adult cohorts and found evidence of same-strain colonization only in seven individuals^70^. On the opposite, if species from the oral cavity are frequently and continuously translocating to the intestines as suggested by Schmidt et al.^7^, the digestive tract would host a connected population of highly similar strains. Our results demonstrated that fecal isolates are more closely related to oral isolates from the same individuals than to isolates from other individuals. This translocation mechanism is consistent with a source-sink dynamic model in which *S. salivarius* from the oral cavity are continuously seeding the lower GI tracts to maintain a connected individual-specific population along the digestive tract. These observations are further in concordance with the findings of Lee et al.^73^ who showed by modeling temporal changes in a longitudinal metagenomic dataset from fecal samples that *Streptococcus* species are transiently colonizing species. Nevertheless, while bacterial isolation consistently detected translocation of *S. salivarius* in the majority of children, metagenomic resolved methods detected a limited number of translocation events per individuals. Based on the nucleotide diversity of *S. salivarius*, several strains are likely to co-exist in each body compartment, as demonstrated by Chen-Liaw et al.^74^ who previously demonstrated in 100 individuals that the strain richness of *S. salivarius* was above two in the feces. The presence of conspecific strains likely affected negatively our ability to reconstruct MAGs and reduced the likelihood to isolate less abundant strains which might have led to gaps in the detection of translocation events^72^.

Although we identified oral-to-gut translocation mechanism, we did not observe a difference in the number of translocation events between stunted and non-stunted children. Indeed, in children suffering from undernutrition, overrepresentation of oral bacteria in the small intestine and the feces has been proposed as a signature of stunting^10^. Several studies taking place in Madagascar, Central African Republic, Pakistan and Bangladesh have shown a decompartmentalization of the GI tract with an overgrowth of oral bacteria within the small intestine^10–12,14^. Therefore, we set out to quantify the total bacterial biomass in the small intestine and feces to assess for the mechanisms leading to an enrichment of oral bacteria in the GI tract. As demonstrated by Liao et al.^22^, the increased relative abundance of oral bacteria in the feces is likely explained by a reduction of the total bacterial load leading to a relatively higher proportion of transient oral bacteria. Using the bacterial-to-host ratio^51^, we demonstrate that the total bacterial load in the feces was positively correlated with the height-for-age z-score. Additionally, in our study, differential abundance analysis shows a potential depletion of butyrate-producing bacteria, consistent with previous observations that highlighted an underrepresentation of these taxa in the feces of undernourished children^10,15^. These results suggest that the enrichment of oral bacteria in the feces of stunted children may be due to the overall reduction of lower GI tract bacteria, contrasting with the mechanism observed in the small intestine. Indeed, based on the total microbial load obtained by qPCR, we found that stunted children in our study suffered from SIOBO, consistent with the results from Vonaesch et al.^11^, who found SIOBO in more than 88% of stunted children from Madagascar and Central African Republic. These results suggest that the overrepresentation of oral bacteria in the small intestine may be due to a change in the duodenal environment allowing oral bacteria to expand. Based on a previous report, children produce an estimated 0.22mL of saliva per minute^75^ containing about 10^6^ CFU/mL of bacteria^76^, resulting in an estimated 3.2*10^8^ CFU/mL bacterial cells swallowed per day. Assuming a reduction by at least 5 orders of magnitude during gastric passage^7,77^, 3.2*10^3^ bacterial cells per day could reach the duodenum. In comparison, the duodenal lumen in children aged 2-5 years contains approximately 40mL of fluid^78^ with a bacterial concentration below 10^5^ CFU/mL for non-SIOBO children and above 10^5^ CFU/mL in those with SIOBO. Consequently, the influx of viable bacteria from daily swallowed saliva would account for only about 0.001% of the duodenal bacterial abundance or up to about 1% using less conservative estimates. The presence of SIOBO in the duodenum of stunted children and the minimal direct contribution of the swallowed bacterial cells suggests that oral bacteria enrichment may be due to a change in the intestinal environment, favoring the growth of oral bacteria in this compartment. Overall, these observations suggest that while the magnitude of translocation from the oral cavity to the lower GI tract does not differ between stunted and non-stunted children, the consequences of oral bacteria consistently seeding the intestines may be more profound in stunted children due to favorable conditions for oral overgrowth in their duodenum and the depletion of colonic bacteria.

Nonetheless, this study is limited to a small number of individuals and larger cohorts sampling multiple body sites are necessary to validate both the translocation and the enrichment mechanisms observed in our study. Furthermore, future studies should include isolates for other bacterial species, such as *Rothia mucilaginosa* and members of the *Veillonella* genera, to generalize our results to commonly transmitted oral bacteria. As our samples were collected at a single time point, future research would benefit from longitudinal profiling of the microbiome, similarly to Zhao and Lieberman et al.^79^. who combined metagenomic and isolated genomes sequencing to demonstrate that *Bacteroides fragilis* undergoes within-host evolution and adaptation for long-term residency. Longitudinal data would help determine whether *S. salivarius* constantly replaces the existing population in the lower GI tract, or whether co-existing strains accumulate mutations, allowing them to thrive in the context of disease.

In summary, by combining metagenomic and isolate-based strain tracking, we provide mechanistic insights by which bacteria of oral origin are translocating from the oral cavity to the lower GI tract. We demonstrated that *S. salivarius* is continuously translocating from the oral cavity to the lower digestive tract following a source-sink dynamic model both in stunted and non-stunted children from Central African Republic. Furthermore, we identified the mechanisms by which oral bacteria are enriched in the small and large intestines of stunted children. Our study underscores the relevance of investigating not only the fecal microbiota but the entire GI tract as a continuum to further understand disease signatures, especially in the context of childhood undernutrition. Identifying translocation and enrichment mechanisms of bacteria of oral origin in the intestinal tract is key to further inform treatment schemes targeting not only the large intestine but all compartments of the digestive tract of stunted children.

## Author contributions (CREediT roles)

Conceptualization: P.V., P.J.S.

Data curation: P.V., S.Y.

Formal analysis: S.Y.

Funding acquisition: P.V., P.J.S.

Investigation: S.Y., Y.T, A.R., E.D., C.N., S.S.V., K.K., N.K., F.M.

Methodology: S.Y., P.V., P.J.S., J.C.G.

Project administration: P.V., S.D., K.K., J.C.G.

Supervision: P.V., S.D., J.C.G., P.J.S.

Visualization: S.Y.

Writing – original draft: S.Y., P.V.

Writing – review & editing: Y.Z., A.R., F.M., J.C.G., E.D., C.N., S.S.V., K.K., S.D., P.J.S., N.K.

## Funding

The Afribiota project received funding through the Total Foundation, Institut Pasteur, the Bill and Melinda Gates Foundation (OPP1204689, INV-004352 and INV-002525), the Fondation Petram and a donation by the Odyssey Re-Insurance company. PV was supported during the Afribiota project by an Early Postdoctoral Fellowship (P2EZP3_152159), an Advanced Postdoctoral Fellowship (P300PA_177876) as well as a Return Grant (P3P3PA_177877). Work in her lab is funded through an Eccellenza Professorial Fellowship (PCEFP3_194545) and a SNSF Starting Grant (TMSGI3_218455) from the Swiss National Science Foundation. This study has been further supported as a part of the NCCR Microbiome, a National Center of Competence and research, funded by the Swiss National Science Foundation (Grant number 180575). The funders had no role in study design, data collection and analysis, decision to publish, or preparation of the manuscript.

## Acknowledgments

We thank all children and their families who participated in the TITI project and further accepted to participate to the present study. Further, we thank the whole Afribiota team including Jean-François Dammaras from Institut Pasteur de Bangui and Dr. Jane Deuve from Institut Pasteur, who administered the financial and logistic aspects of the project, the Complexe Pédiatrique de Bangui, the Institut Pasteur de Bangui, the Institut Pasteur, the Hospital Necker-Enfants malades,the Hopital Pitié-Salpêtrière and the University of Lausanne. We also thank Myriam Corthesy for her help with the MALDI-TOF spectroscopy measurements. We thank Dr. Naika Prince for her help setting up the qPCR, Dr. Garance Sarton-Lohéac for her precious advices on phylogenetic tree building and Malick Ndiaye for his help and advice on the metagenomic analysis. We further thank the whole Vonaesch lab, Prof. Philipp Engel as well as several colleagues at the Department of Fundamental Microbioloy, University of Lausanne, for fruitful discussions. Chat GPT-5 was used for language improvement and proof reading.

## Declaration of interest statement

The authors declare no competing interest.

## Appendix A

The AFRIBIOTA Investigators who co-auhored this article are composed as follows (in alphabetical order):

Jean-Marc Collard, Institut Pasteur de Madagascar, Antananarivo, Madagascar Maria

Doria, Institut Pasteur, Paris, France

Darragh Duffy, Institut Pasteur, Paris, France

Serge Ghislain Djorie, Institut Pasteur de Bangui, Bangui, Central African Republic

Tamara Giles-Vernick, Institut Pasteur, Paris, France

Bolmbaye Privat Gondje, Complexe Pédiatrique de Bangui, Bangui, Central African Republic

Jean-Chrysostome Gody, Complexe Pédiatrique de Bangui, Bangui, Central African Republic

Milena Hasan, Institut Pasteur, France

Nathalie Kapel, Hôpital Pitié-Salpêtrière, Paris, France

Jean-Pierre Lombart, Institut Pasteur de Bangui, Bangui, Central African Republic

Synthia Nazita Nigatoloum, Complexe Pédiatrique de Bangui, Bangui, Central African Republic

Laura Wegener Parfrey, University of British Columbia, Vancouver, Canada

Maheninasy Rakotondrainipiana, Institut Pasteur de Madagascar, Antananarivo, Madagascar

Pierre-Alain Rubbo, Institut Pasteur de Bangui, Bangui, République Centrafricaine

Philippe Sansonetti, Institut Pasteur, Paris, France

Ionela Gouandjika-Vasilache, Institut Pasteur de Bangui, Bangui, République Centrafricaine

Pascale Vonaesch, Institut Pasteur, Paris, France

Sonia Sandrine Vondo, Complexe Pédiatrique de Bangui, Bangui, Central African Republic

---

**Supplementary Figure 1:**
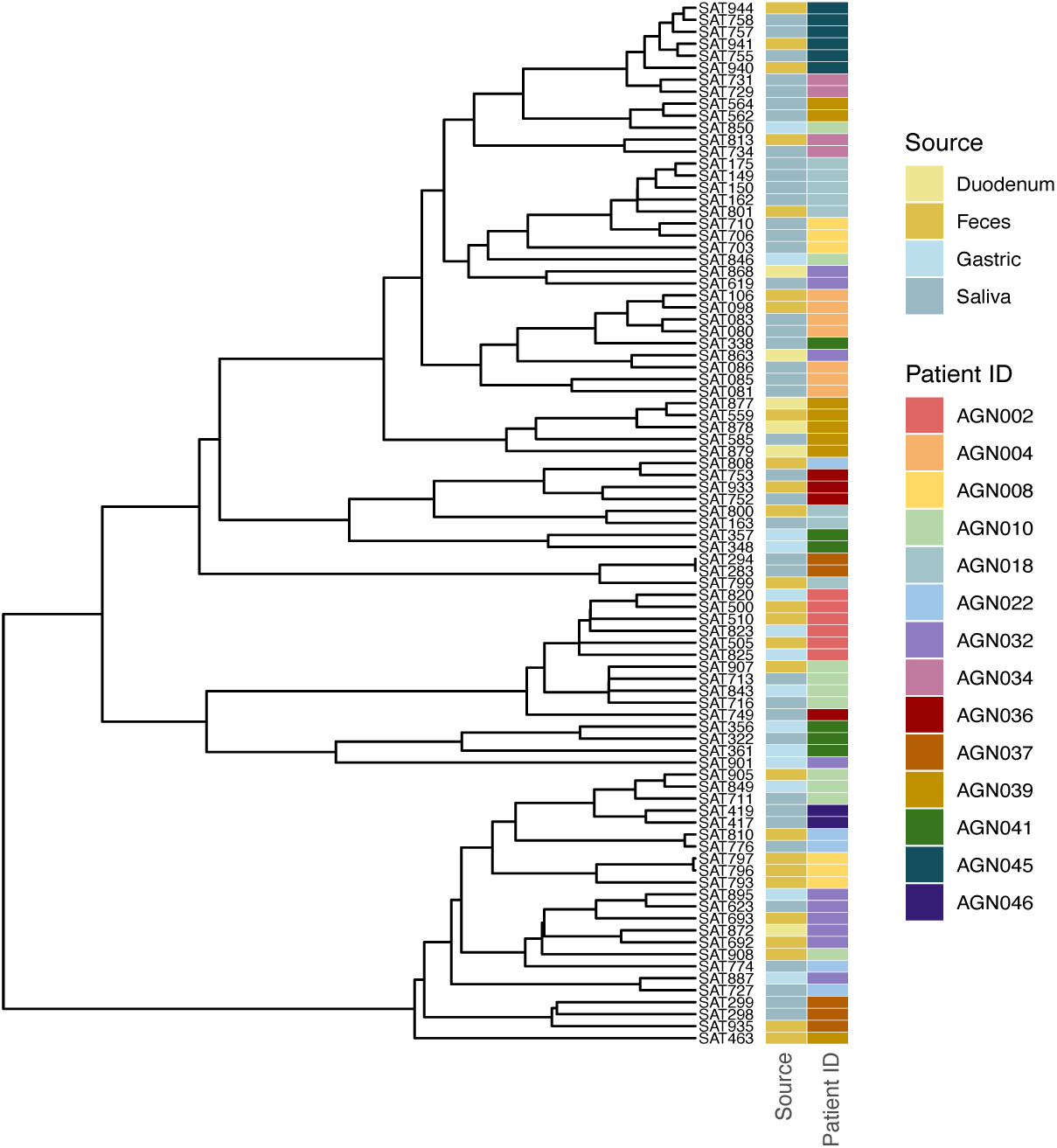
Ultrametric phylogeneic tree of *S. saliavrius* genomes of all isolates gathered in this study.

## Notes

### Competing Interest Statement

The authors have declared no competing interest.

